# Body Maps of Uncertainty and Surprise in Musical Chord Progression and its individual differences in Depression and Body Perception Sensitivity

**DOI:** 10.1101/2023.09.07.556777

**Authors:** Tatsuya Daikoku, Masaki Tanaka, Shigeto Yamawaki

**Author notes:** **Corresponding author** Tatsuya Daikoku, Graduate School of Information Science and Technology, The University of Tokyo, Tokyo, Japan, 7-3-1 Hongo, Bunkyo-ku, 113-8656, Tokyo, Japan. **Competing Interests** The authors declare no competing financial interests. **Author Contributions** T.D. and M.T. conceived the experimental paradigm and method of data analysis. T.D. collected the data, analysed the data, and wrote the draft of the manuscript and figure. T.D., M.T. and S.Y. edited and finalized the manuscript. **Data Availability** All of anonymized raw data files, stimuli used in this study and the results of statistical analysis have been deposited to an external source (https://osf.io/cyqhd/). The other data are shown in supplementary data.

## Abstract

Music has profoundly shaped the human experience across diverse cultures and generations, yet the mechanisms that it influences our minds and bodies remain elusive. This study examined how the perception of music chords elicits bodily sensations and emotions through predictive processing of the brain. By deploying body-mapping tests and emotional evaluations on 527 participants exposed to chord progressions, we unveil the intricate interplay between musical uncertainty, prediction error, and temporal dynamics in eliciting specific bodily sensations and emotions. Our results demonstrated that the chord progressions characterized by low uncertainty coupled with high surprise or predictability evoke bodily sensations closely associated with interoception including the cardiac and abdominal regions. Notably, these sensations are associated with aesthetic appreciation, with the intensity of cardiac sensations being positively correlated with valence in chord progressions with low uncertainty and high surprise. These results highlight the pivotal role of uncertainty and prediction error in shaping emotional responses and also suggest a hypothesis for emotion generation through predictive processing and sound embodiment. This study offers a tantalizing glimpse into the potential nexus between interoception by music and mental well-being, underscoring the importance of recognizing diverse forms of musical pleasure and their unique effects on our minds and bodies.

## 1. Introduction

Music, an omnipresent force throughout human history, has deeply influenced our minds and bodies (Mithen, 2006). Its pervasive and far-reaching effects have captivated not only the scientific community but also a broad audience, spanning diverse cultures and generations. Despite centuries of inquiry into the myriad ways in which music influences our minds and bodies, this intriguing question remains largely unanswered and offers fertile ground for new discoveries and insights.

A promising avenue to unlock this enigma revolves around the unique bodily sensations elicited by music (Putkinen et al., 2023; Hove et al., 2020). Beyond the external sensory perceptions (i.e., exteroception) like an auditory system music elicits internal bodily perceptions such as interoception (e.g., heartbeat acceleration or the spine-chilling thrill) and proprioception (e.g., the constrictive sensation in one’s chest when hearing sorrowful music) (Trost et al., 2017; Mori et al., 2017; Koelsch, Jancke, 2015). Particularly, interoceptive awareness, driven by shifts in interoceptive sensations, has been identified as being integral to our mental (Critchley et al., 2004) and potentially associated with music emotion and its embodiment (Christensen et al., 2018; Schirmer-Mokwa et al., 2015). Although research has richly explored how music hearing through exteroception influences our emotions, the intricate interplay between music and internal bodily perceptions involved in emotions remains largely unexplored.

The emotions are tightly interwoven with our bodily sensation. A past study revealed that the different emotions can be discerned through the mapping of emotion-triggered bodily sensations (Nummenmaa et al., 2014). They suggested that diverse emotions have distinct body topographies with negative emotions such as fear, anger, sadness, and anxiety activating the upper side of the body and positive ones such as happiness and love casting a wider range of activation. This implies that emotions are embodied, manifesting spatially within the body. Further, the mappings of emotion-triggered bodily sensation may serve as a key to understanding individual differences in interoceptive awareness. For example, persons who exhibit weak interoceptive awareness, including individuals with depression and alexithymia (Herbert et al., 2011; Ernst et al., 2014; Pollatos et al., 2009), show weak or diffused emotion-triggered bodily sensations in individuals with depression (Lyons et al., 2021; Lloyd et al., 2021). Further, a past study revealed interoceptive accuracy was associated with emotion-triggered bodily sensations (Jung, Ryu et al., 2017). They suggest that the awareness of one’s internal bodily states might play a crucial role as a required messenger of sensory information during the affective process.

The relationship between bodily sensations and emotions can be elucidated from the perspective of the brain’s predictive processing. The brain generates emotions by minimizing prediction errors between the anticipatory signals derived from its internal model and the sensory signals through exteroceptive, interoceptive, and proprioceptive sensations (Seth et al., 2013). It is suggested that interoceptive awareness arises from the updating of prediction errors (Barrett et al., 2015; Khalsa et al., 2015; Ainley et al., 2016). For example, if a musical chord progression ends suddenly with an unexpected shift to a different key, leading to a sudden increase in interoceptive prediction error (e.g., heartbeat accentuation), then the consequent interoceptive prediction error may be resolved by updating interoceptive priors, possibly through deep breathing or other measures.

Regarding music prediction and the elicited emotion, recent research has illuminated the critical interplay of both predictive uncertainty and surprise (predictive error) (Cheung et al., 2019; Huron, 2008; Daikoku et al., 2021). In a study by Cheung et al. (2019), they developed a predictive model that mathematically decodes the relationship between uncertainty and surprise in music. Analyzing 80,000 chords from US Billboard pop songs, they anchored their study in the framework of Western tonal harmony, viewing it as a representation of musical syntax. The study unearthed that chords combining low uncertainty with high surprise or those combining high uncertainty with low surprise were the most pleasurable. Neurologically, this effect correlated with activity in regions such as the amygdala, hippocampus, and auditory cortex. Notably, dopaminergic pathways in the nucleus accumbens, which responded positively to uncertainty, seemed to play a pivotal role. Their findings echoed Berlyne’s seminal model, where pleasure forms an inverted-U relationship with factors like complexity. However, the multifaceted nature of musical pleasure, as Cheung et al. illustrated, suggests that our musical experiences are shaped by a web of interacting variables such as uncertainty-weighted inverted U relationship, rather than isolated factors. In recent decades, the relationship between exteroceptive audition (auditory cortex), and music emotion based on predictive processing is progressively being elucidated. However, the associations of music emotion with interoception and bodily sensations represented as the body maps continue to be an under-researched frontier.

This study embarks on this unexplored voyage. We aspire to elucidate how perceptions of musical chord progressions are embodied, giving rise to emotional experience, grounded in the framework of the body map of emotions. Specifically, we aim to determine what types of musical chords, through the lens of predictive processing, evoke interoceptive bodily sensations, particularly those related to the cardiac and abdominal (stomach) regions. These two regions are particularly known to be associated with interoceptive awareness and interoceptive prediction error (Critchley and Harrison, 2013; Seth, 2013; Garfinkel et al., 2015). Rooted in predictive processing principles, our central hypothesis is that two factors: uncertainty-weighted prediction error and its fluctuations (temporal dynamics) critically contribute to interoception evoking bodily sensations.

With respect to the element of uncertainty-weighted prediction error, based on the uncertainty-weighted inverted U relationship characterizing musical pleasure, we posited that chords combining low uncertainty with high surprise or those combining high uncertainty with low surprise induce the interoception evoking bodily sensations. Notably, it has been known that the increasing complexity (uncertainty) of regularities requires an increasing amount of knowledge about musical regularities to make precise predictions about upcoming musical events (Rohrmeier and Rebuschat, 2012) and that the correct and precise predictions may be perceived as more rewarding (Koelsch, 2015). Consequently, for the general people who are not experts in music, chords combining low uncertainty with high surprise might be particularly salient in accurately recognizing prediction errors, thereby potentially amplifying interoceptive bodily sensations and the associated emotions (Singer, Critchley et al., 2009).

Concerning the element of the fluctuations (temporal dynamics) of uncertainty and surprise, the facilitation of interoceptive bodily sensations by uncertainty-weighted prediction error could be modulated by preceding musical contexts. That is, while it has been known revealed that the musical pleasure forms a two-dimensional inverted U curve based on uncertainty and surprise, music entails extended contexts that cannot be fully captured by the simple prediction and uncertainty between one chord and the next. The music context can influence the perception of even the same two-chord sequence, resulting in different emotional experiences depending on the surrounding context. Consequently, in addition to the two dimentions of uncertainty and surprise, it is necessary to elucidate musical prediction and emotion through a three-dimensional model that takes into account the preceding contexts (more dynamics). These temporal dynamics of uncertainty-weighted prediction error could offer a more nuanced understanding of the emotions and bodily sensations intrinsic to music as a temporal art form.

## 2. Results

All participants (*N* = 527) conducted body-mapping tests and the following emotional judgements in every eight types of 4-chord progression (see the Methods section for details). These chord progressions were generated using a statistical-learning model (Daikoku, Minaotya et al., 2023) to compute the Shannon information content and entropy, based on transitional probabilities (Shannon, 1948) of each chord using a corpus of 890 pop songs from the US Billboard (Burgoyne, Wild, and Fujinaga, 2011). Entropy gauges the perceptual uncertainty a listener feels in predicting an “upcoming” chord based on prior chords, while information content quantifies the surprise experienced upon hearing the actual chord.

Using this model, we generated the 92 unique chord progressions encompassed within the eight types of chord progressions. Each type is characterized by varying degrees of uncertainty and surprise (see red and blue lines in Figure 1). Four of the 8 types began with three chords, each with low surprise and uncertainty (SluL: a-d in Figure 1), while the other four began with three chords each displaying high surprise and uncertainty (sHuH: e-h in Figure 1). For each chord progression, the last (i.e., 4th) chord was generated with a 2×2 pattern, varying in uncertainty and surprise. The first pattern exhibited both low surprise and low uncertainty (sLuL: Figure 1a and 1e), the second had low uncertainty but high surprise (sHuH: Figure 1b and 1f), the third showcased low surprise with high uncertainty (sLuH: Figure 1c and 1g), and the fourth possessed both high surprise and uncertainty (sHuH: Figure 1d and 1h). The thresholds for the high and low values were established based on the top and bottom 20% of all data points for each uncertainty and surprise. Multiple chord progressions were generated for each eight types, and the chord progression employed was randomly selected for each participant.

**Figure 1.**
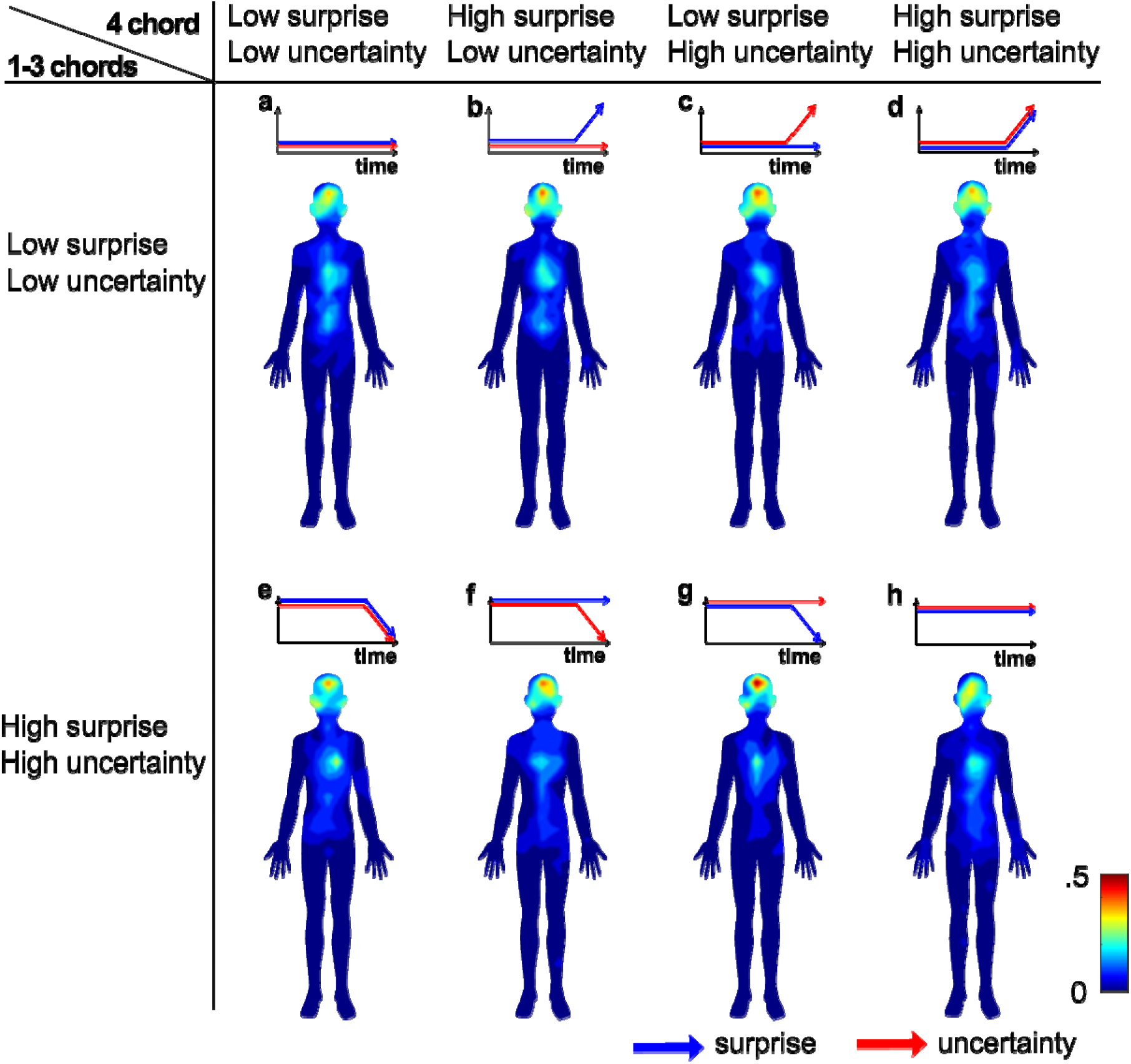
Body topography of musical chord progressions. The blue-to-red gradients represent the number of clicks. The blue and red arrows represent the surprise and uncertainty values, respectively.

Participants were exposed to these eight types of chord progressions in random order. Following each listening session, they were asked to respond with clicks to the position in the body where they felt from the chords, using the body image presented on the screen. Two types of emotional judgements were administered. The first comprised multiple-choice categorical judgements; that is, in each type of chord progression, participants were required to select the best 5 emotional categories in ranking elicited by each sound from a list of 33 categories. The second kind comprised nine-point Likert scales of valence and arousal. We compared the topography of chords and the corresponding emotional responses among the eight types of chord progressions.

### 2.1. Body map

Grand averages of the body map and the clicked positions were shown in Figure 1 and supplementary materials, respectively. All the anonymized raw data files and all the results of statistical analyses including the descriptive have been deposited to an external source (https://osf.io/cyqhd/). As illustrated in Figure 1, each chord progression revealed a distinct body map. Particularly, a specific type of chord progression transitioning from predictable 1st-3th chords to the last (4th) chord characterized by low uncertainty and high surprise (Figure 1b) provoked pronounced bodily sensations localized to the heart (see Figure D in the Supplementary). Further, a chord progression characterized by only predictable chords (Figure 1a) provoked pronounced bodily sensations localized to the abdomen.

The Shapiro–Wilk test for normality showed the violations of the assumption of normality on all the data (p < .001). Hence, we applied Friedman’s non-parametric Repeated-measure analysis of variance (ANOVA) for within-subject factor “types of chord progressions”. The clicks at the abdomen position showed significant main effects (χ*² =* 16.7, df = 7, *p =* .019). The Durbin-Conover post hoc test detected that the abdomen sensation was stronger in the predictable chord progressions with low uncertainty (Figure 1a) than in the chord progressions transitioning from unpredictable chords to the last chord characterized by high uncertainty and low surprise (Figure 1g) (p<.001).

### 2.2. Emotion in response to musical chord progressions

All the results of statistical analyses and the descriptive of the multiple-choice categorical judgements and nine-point Likert scale of valence and arousal have been deposited to an external source (https://osf.io/cyqhd/). The figures of the grand average data are shown in the Supplementary (Figure C and E). The Shapiro–Wilk test for normality showed the violations of the assumption of normality on valence, arousal, and aesthetic ratings (p < .001). Hence, we applied the Friedman’s non-parametric repeated-measure ANOVA for within-subject factor: 8 types of chord progressions.

The ANOVA for valence showed the main effects (χ*² =* 215, *p <* .001). The Durbin-Conover post hoc test detected that the valence was significantly positive in the predictable chord progressions with low uncertainty (Figure 2a, Valence) and the chord progressions transitioning from predictable chords to the last chord characterized by low uncertainty and high surprise (Figure 2b, Valence) compared to the other types of chord progressions (Figure 2c-h, Valence) (p < .001). The valence was significantly positive in the predictable chord progressions with low uncertainty (Figure 2a, Valence) than the chord progressions transitioning from predictable chords to those characterized by low uncertainty and high surprise (Figure 2b, Valence) (p=0.02).

**Figure 2.**
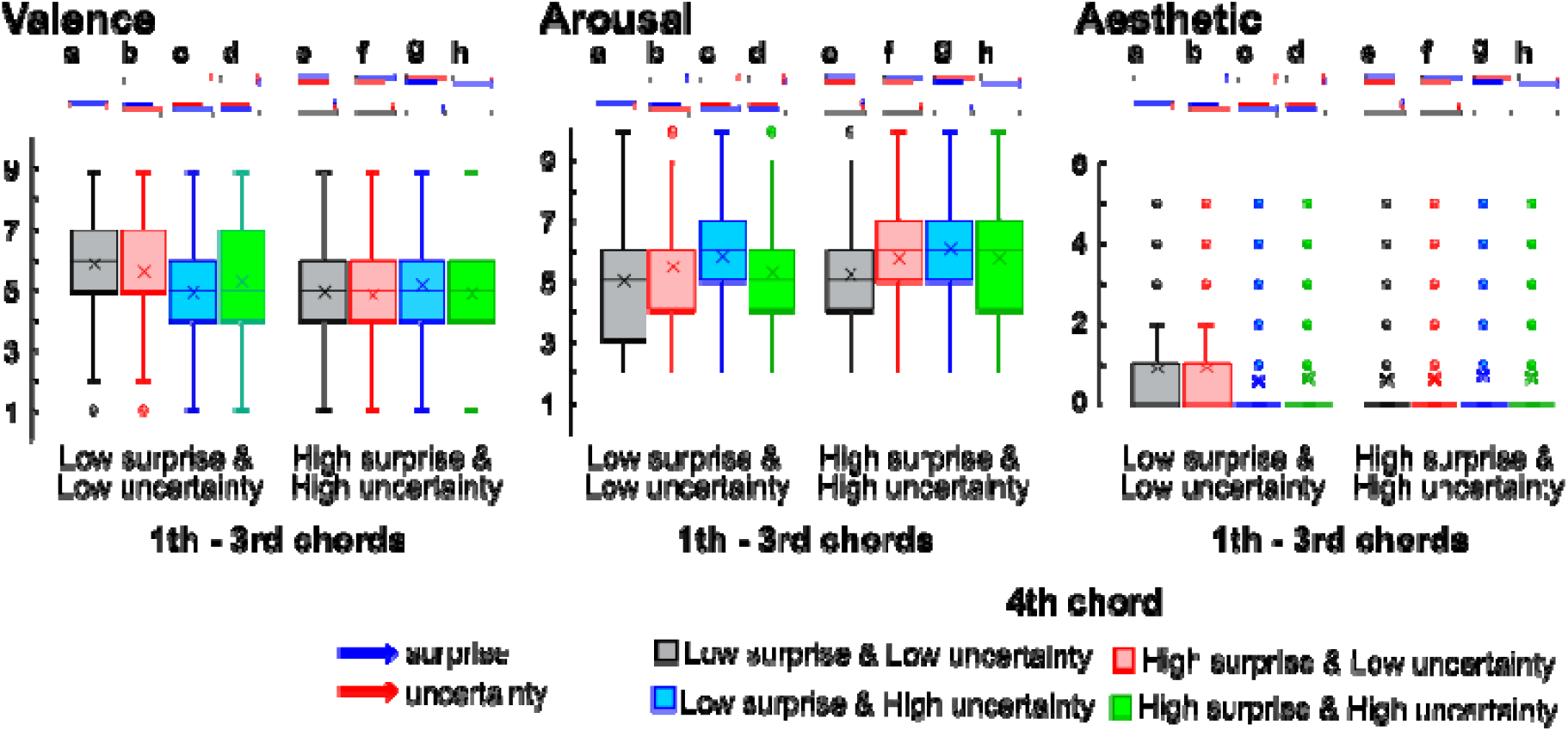
Valence, arousal, and aesthetic ratings for musical chord progressions. The blue and red arrows represent the surprise and uncertainty values, respectively.

The ANOVA for arousals showed the main effects (χ*² =* 192, *p <* .001). The Durbin-Conover post hoc test detected that the arousal was significantly lower in the predictable chord progressions with low uncertainty (Figure 2a, Arousal) and the chord progressions transitioning from predictable chords to the last chord characterized by low uncertainty and high surprise (Figure 2b, Arousal) than the other types of chord progressions (p < .05).

The ANOVA for aesthetic rating showed the main effects (χ*² =* 27.2, *p <* .001). The Durbin-Conover post hoc test detected that the aesthetic rating was significantly higher in the predictable chord progressions with low uncertainty (Figure 2a, Aesthetic) and the chord progressions transitioning from predictable chords to the last chord characterized by low uncertainty and high surprise (Figure 2b, Aesthetic) than the other chord progressions (p < .01).

### 2.3. Correlation between bodily sensation and the emotions

We examined how the total number of clicks at cardiac and abdomen positions were correlated with the scores of valences, arousals, and aesthetic appreciation. The Shapiro–Wilk test for normality showed the violations of the assumption of normality on all the data (p < .001). Hence, we applied the Spearman correlation tests. All the results of statistical analyses and the descriptive have been deposited to an external source. All the figures are shown in Supplementary (Figure F). Results revealed significant positive correlations of valence with the number of clicks localized to the cardiac area only in the chord progressions transitioning from predictable chords to those characterized by low uncertainty and high surprise (Figure 3b) (*rs* = 0.132, *p* = <.001). This suggests that cardiac sensation is an important factor for positive emotion in this type of chord progression.

**Figure 3.**
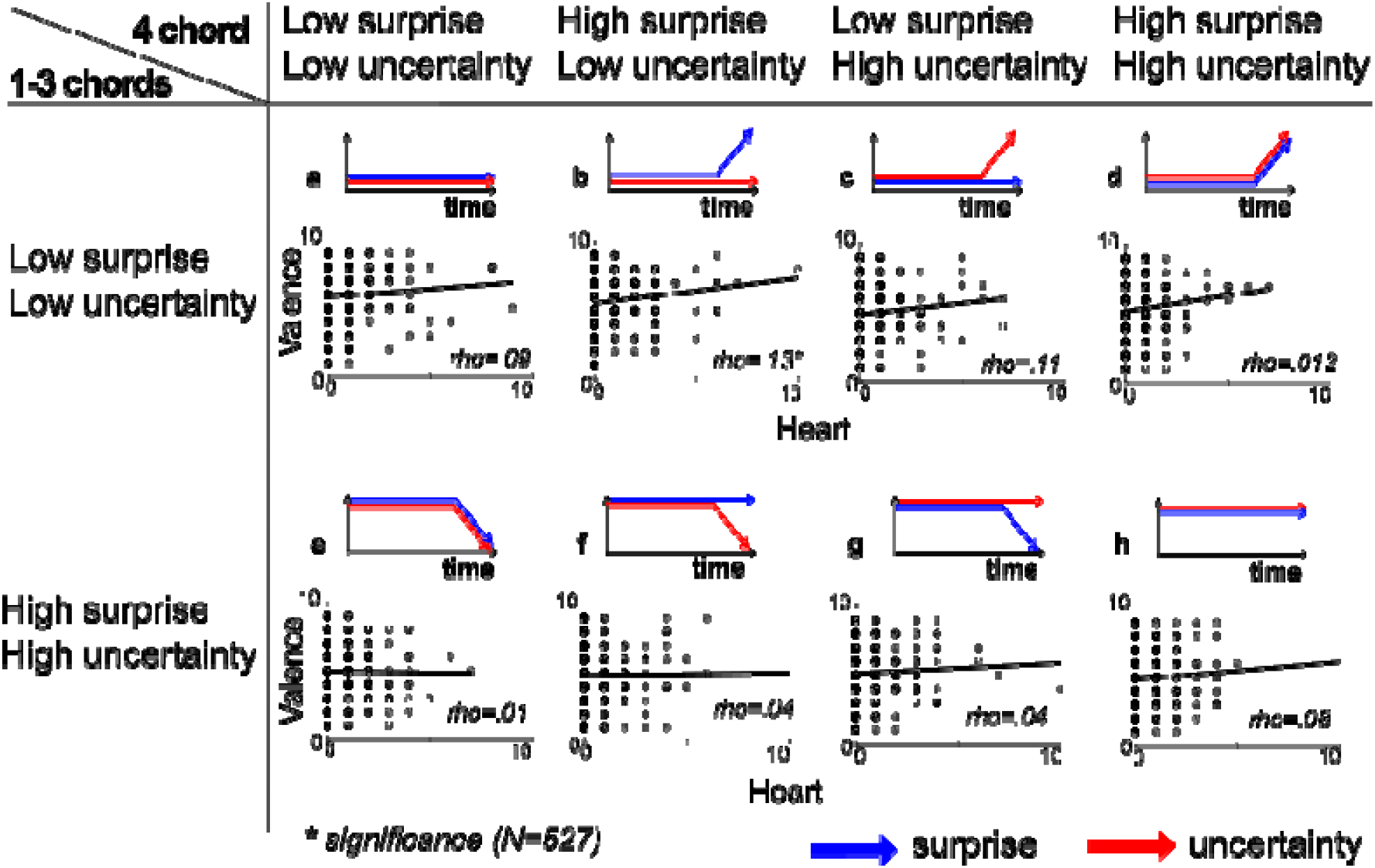
Correlation of valence rating with bodily sensations at cardiac and abdomen areas. The blue and red arrows represent the surprise and uncertainty values respectively in each 8 types of chord progressions.

### 2.4. Emotional Distribution in response to each musical chord progression

To visualize the difference in the emotional distributions between the different types of chord progressions, between groups who rated high and low valence, and between groups who showed strong and weak sensations of each cardiac and abdomen area, we compared the scores of the emotional distributions of 33 emotional categories by Uniform Manifold Approximation and Projection (UMAP) (McInnes et al., 2018).

The 33 emotional categories were separately averaged for each of the 92 distinct chord progressions included within the eight types, as well as for groups with valence scores above 6 and below 4 (Figure 4, top). Additionally, the 33 emotional categories were separately averaged for those who experienced bodily sensations with a click count greater than 1 in the cardiac area and for those who did not experience bodily sensations with a click count of 0 in the cardiac area (Figure 4, bottom left). Lastly, the 33 emotional categories were separately averaged for those who experienced bodily sensations with a click count greater than 1 in the abdominal area and for those who did not experience bodily sensations with a click count of 0 in the abdominal area (Figure 4, bottom right).

**Figure 4.**
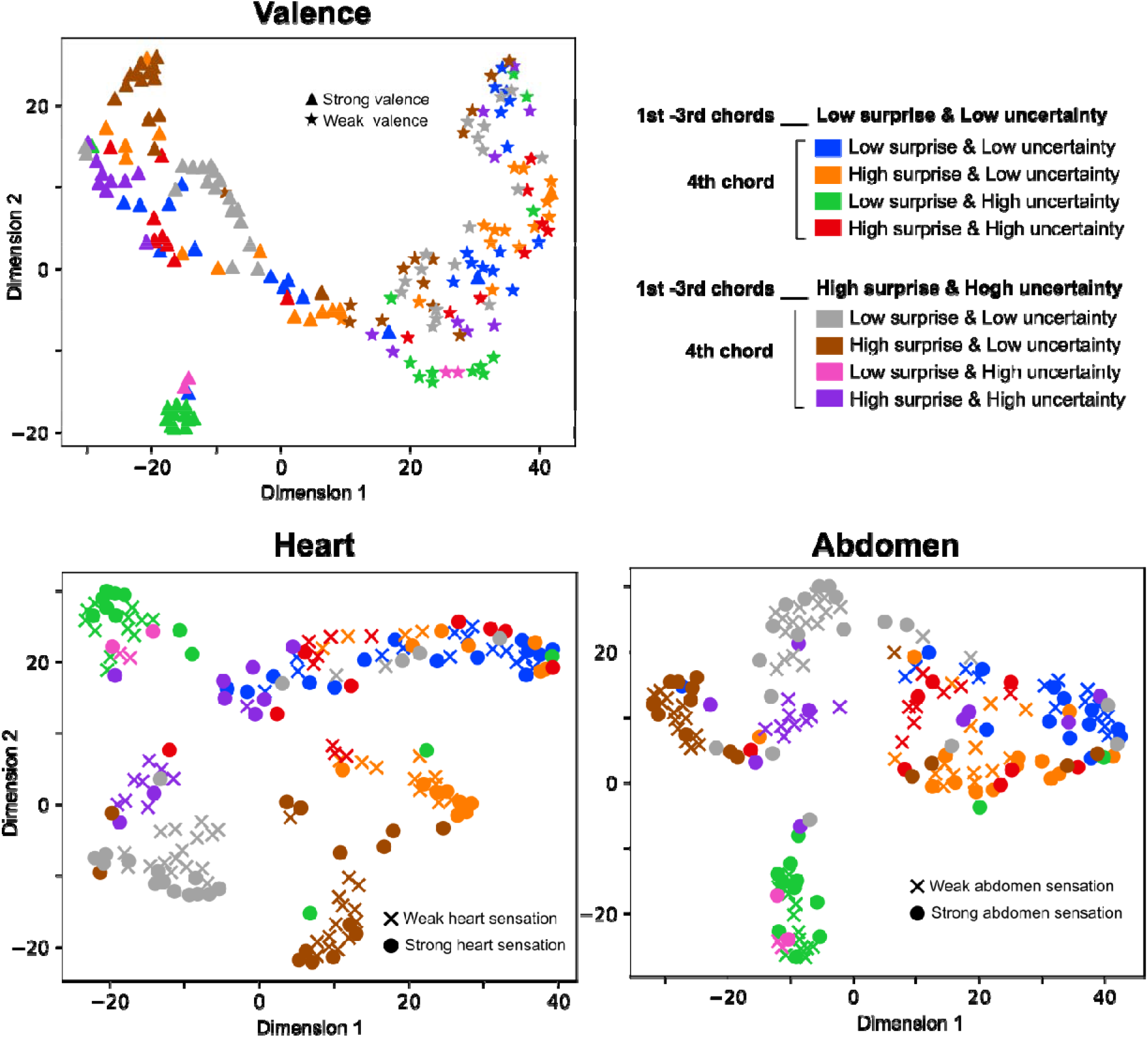
The structure of reported emotional experience. A chromatic map of average emotional responses to 92 chord progressions within a 33-dimensional categorical space of reported emotional experience. Uniform Manifold Approximation and Projection (UMAP) was applied to the loadings of the 92 chord progressions on the 33 categorical judgment dimensions, generating loadings of each stimulus on two axes.

Then, using the averaged data from these 33 emotional categories, we employed UMAP in Python to craft a two-dimensional representation. The results showed that the intricate interplay between musical uncertainty, prediction error, and temporal dynamics underlies distinct emotional components. Notably, even when different types of chord progressions culminate in a singular positive valence, the dynamics of musical chords, especially those transitioning from predictable chords to those characterized by high uncertainty and low surprise, manifest uniquely.

## 3. Discussion

The present study examined how perceptions of musical chord progressions elicit bodily sensations and emotions. Specifically, within the framework of predictive processing, we aim to determine how chord prediction and its uncertainty evoke interoceptive bodily sensations, particularly around the cardiac and abdominal regions. Our findings indicate that the chord progressions characterized by low uncertainty and high surprise as well as the predictable chord progressions with low uncertainty (Figure 1b and 1a) provoked cardiac and abdomen bodily sensations, respectively. Further, such bodily sensations were correlated with an aesthetic appreciation (Figure 2). This effect was pronounced when the preceding musical context comprised predictable and less uncertain chords (Figure 1a-d), in contrast to sequences typified by high prediction errors and high uncertainty (Figure 1e-h). Notably, in the chord progressions characterized by low uncertainty and high surprise, the intensity of cardiac sensations was positively correlated with valence (Figure 3). Our findings also suggest that the different interplay of uncertainty and prediction error elicits distinct emotional distributions, especially in a sequence transitioning from predictable chords to a chord with high uncertainty and low surprise, even when all these chord progressions converge on a positive valence.

Our past study investigating the body map of sound pitch revealed that the lowest bodily location in response to pitch was not the feet, located at the lowest point of the body, but rather the abdomen, the lowest part of the visceral system (Daikoku, Horii, Yamawaki, 2023, preprint). Further, persons who tend to show weak interoceptive awareness such as alexithymia and depression exhibited less localized and more diffuse bodily sensations in response to pitch. Such diffused bodily sensation is correlated with strong feelings of anxiety and negative valence in response to several pitches, suggesting that the diffuse bodily sensations may induce negative emotions such as anxiety. Thus, emotional experience induced by auditory perception has been suggested to involve the ‘embodiment’ of sound through proprioceptive and interoceptive pathways. In this study, by specifically employing musical stimuli of chords instead of simple pitches, we more definitively highlighted interoceptive bodily sensations including the cardiac (heart) and abdominal (stomach) regions.

A past study detected that chords combining low uncertainty with high surprise or those combining high uncertainty with low surprise were the most pleasurable (Cheung et al., 2019). This suggests that the musical pleasure forms a two-dimensional inverted U curve based on uncertainty and surprise. However, music entails extended contexts that cannot be fully captured by the simple prediction and uncertainty between one chord and the next. This context can influence the perception of even the same two-chord sequence, resulting in different emotional experiences depending on the surrounding context. Consequently, in addition to the two dimentions of uncertainty and surprise, it is necessary to elucidate musical prediction and emotion through a three-dimensional model that takes into account the preceding contexts (more dynamics). The present study first revealed that the emotional responses induced by uncertainty-weighted prediction error could be modulated by preceding musical contexts. In other words, the temporal dynamics of uncertainty-weighted prediction error could offer a more nuanced understanding of musical emotions as a temporal art form.

We also found that the three-dimensional interplay between musical uncertainty, prediction error, and temporal dynamics underlies distinct emotional components. Interestingly, even when different types of chord progressions culminate in a singular positive valence, the different temporal dynamics (fluctuation) of surprise and uncertainty manifest uniquely. It’s worth acknowledging the diverse forms of musical pleasure and distinguishing between these could elucidate distinct effects on our mind and body.

This study proposes a hypothesis for emotion generation from auditory perception, emphasizing the embodiment of sound through predictive processing. That is, we postulate that positive emotions emerge during the process of sound embodiment, wherein auditory bodily sensations become localized, i.e., prediction errors are minimized, through proprioceptive and interoceptive predictive processing. Through computational modelling, a past study indicated that individuals exhibiting hypo-sensitivity to stimuli, as seen in depression or alexithymia (Dunn et al., 2007; Ehlers et al., 1996; Serafini et al., 2016), might struggle to adapt to auditory sensory signals (Daikoku, Minaotya, Kamamerns, 2023). Consequently, the internal model’s updating mechanism, aimed at minimizing prediction errors, may become dysfunctional. Given these findings, the observed the less localized body maps and amplified anxiety in prior research on alexithymia and depression (Daikoku, Horii, Yamawaki, preprint) may be attributed to a malfunctioning error-minimization mechanism due to hypo-sensitivity. This malfunction might subsequently lead to emotions not being correctly identified, giving rise to anxiety.

Beyond the exteroception such as hearing, music induces feelings inside the body such as interoception and proprioception. Particularly, interoceptive awareness has been identified as being integral to our mental well-being (Critchley et al., 2004). Thus, profound interoceptive changes induced by music might potentiate interoceptive awareness, thereby potentially benefiting mental health.

This study did not directly investigate whether interoceptive awareness was enhanced by the perception of musical chords. In other words, it is possible that bodily sensations were simply induced in the chest and abdominal areas regardless of interoception. Future research should investigate interoception such as a heartbeat detection test before and after music listening to determine whether interoceptive awareness is enhanced. Furthermore, it is of note that the strongest bodily sensations were generated around the head and ears. This effect is clearly due to exteroception (i.e., hearing). It is necessary to elucidate the fundamental differences between exteroceptive and interoceptive body maps in future research.

Nevertheless, this study is the first to demonstrate that emotional responses to music based on predictive processing can induce specific bodily sensations. Particularly, even when different types of chord progressions culminate in a single positive valence, the varying temporal dynamics of surprise and uncertainty manifest uniquely. It is important to acknowledge the diverse forms of musical pleasure, and distinguishing between these could shed light on their distinct effects on our minds and body.

## 4. Conclusions

This study reveals the intricate interplay between musical uncertainty, prediction error, and temporal dynamics in eliciting distinct bodily sensations and emotions. We found that specific types of chord progressions provoke sensations in the cardiac and abdominal regions. These sensations are associated with aesthetic appreciation. Furthermore, our findings underscore the importance of acknowledging diverse forms of musical pleasure, each with unique effects on our minds and bodies. We propose a novel hypothesis for emotion generation through predictive processing and sound embodiment, suggesting a potential link between musical interoception and mental well-being. Further research is required to explore this connection and to differentiate between exteroceptive and interoceptive bodily sensations in music perception.

## 4. Methods

### 4.1. Participants

The present study consists of body-mapping tests and the following emotional judgements in every eight types of 4-chord progression. The Japanese participants took part in the study (N = 527, *M*_age_±SD = 32.18_±_5.37, female = 257, specific musical training_±_SD = 2.47_±_5.66). They have no history of neurological or audiological disorders and no absolute pitch. The experiment was conducted in accordance with the guidelines of the Declaration of Helsinki and was approved by the Ethics Committee of The University of Tokyo (Screening number: 21-335). All participants gave their informed consent and conducted the experiments by PC.

### 4.2. Materials and Methods

The experimental paradigm was generated using Gorilla Experiment Builder (https://gorilla.sc), which is a cloud-based research platform that allows deploying of behavioural experiments online. Each participant was provided with eight types of chord progression including 4 chords (500ms/chord, 44.1kHz, 32bit, Electric Piano 1 based on, General MIDI, amplitude based on equal-loudness-level contours).

We used a statistical-learning model (Daikoku, Minaotya et al., 2023) to derive the surprise and uncertainty of every chord. This model computes the Shannon information content and entropy based on transitional probabilities (Shannon, 1948) of each chord from a corpus of 890 pop songs in the McGill Billboard Corpus (Burgoyne, Wild, and Fujinaga, 2011). The statistical learning is an implicit process by which the brain automatically computes transitional probability from sequences (Furl et al., 2011; Yumoto and Daikoku, 2016, 2018), grasps uncertainty/entropy (Hasson, 2017), and predicts a future state based on the internalized statistical model. The transitional probability is a conditional probability of an event en+1, given the preceding n events based on Bayes’ theorem (P(e_n+1_|e_n_)). From a psychological perspective, the transitional probability (P(e_n+1_|e_n_)) can be interpreted as positing that the brain predicts a subsequent event e_n+1_ based on the preceding events e_n_ in a sequence. In other words, learners expect the event with the highest transitional probability based on the latest n states, whereas they are likely to be surprised by an event with a lower transitional probability. Furthermore, transitional probabilities are often translated as information contents:

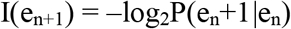

The lower information content (i.e., higher transitional probability) means higher predictabilities and smaller surprising, whereas the higher information content (i.e., lower transitional probability) means lower predictabilities and larger surprising. In the end, a tone with lower information content may be one that a composer is more likely to predict and choose as the next event, compared to tones with higher information content. The information content can be used in computational studies of music to discuss psychological phenomena involved in prediction and statistical learning. the entropy of chord e_n+1_ is the expected information content of chord e_n_. This is obtained by multiplying the conditional probability of all possible chords in S by their information contents and then summing them together, giving:

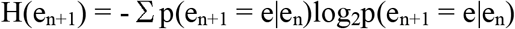

Entropy gauges the perceptual uncertainty a listener feels in predicting an upcoming chord based on prior chords, while information content quantifies the surprise experienced upon hearing the actual chord. Using this model, we generated the 92 unique chord progressions encompassed within the eight types of chord progressions. All the chord progressions have been deposited to an external source (https://osf.io/cyqhd/). Each type is characterized by varying degrees of uncertainty and surprise (see red and blue lines in Figure 1). Four of the 8 types began with three chords, each with low uncertainty and surprise (Figure 1a-d), while the other four began with three chords each displaying high uncertainty and surprise (Figure 1e-h). For each set of these four progressions, the fourth chord was generated with a 2×2 pattern, varying in uncertainty and surprise. Specifically, there are four variations for the fourth chord: the first exhibited both low uncertainty and surprise (Figure 1a and 1e), the second had low uncertainty but high surprise (Figure 1b and 1f), the third showcased high uncertainty with low surprise (Figure 1c and 1g), and the fourth possessed both high uncertainty and surprise (Figure 1d and 1h). The thresholds for the high and low values were established based on the top and bottom 20% of all data points for both uncertainty and surprise. Multiple chord progressions were generated for each eight types, and the chord progression employed was randomly selected for each participant.

Participants were exposed to these eight types of chord progressions in random order. Following each listening session, they were asked to respond with clicks to the position in the body where they felt from the chords, using the body image presented on the screen. The clicking was allowed any number of times up to a maximum of 100 clicks, and clicking while listening to the sound was also allowed (see Figure A in the Supplementary for the details). Two surveys were used to obtain emotional judgements. The first comprised multiple-choice categorical judgements; that is, in each type of chord progression, participants were required to select the best 5 emotional categories in ranking elicited by each sound from a list of 33 categories (see Table in the Supplementary). The 33 emotion categories were derived from emotion taxonomies of prominent theorists, Keltner and Lerner (2010) and based on the previous study by Cowen et al., (2017; 2019; 2020). The second kind comprised nine-point dimensional judgments; that is, after hearing the chord progression, participants were required to rate each type of chord progression along the valence and arousal. Each rating was obtained on a nine-point Likert scale with the number 5 anchored at neutral.

### 4.3. Statistical Analysis

Using the coordinate data of x and y in the body mapping test, we extracted the total number of clicks at two interoceptive positions including cardiac and abdomen areas in each participant. The raw data of x and y coordinates (see Figure B in Supplementary) were down-sampled by a factor of 40. The Figures of the body topographies (Figure 1) were generated using Matlab (2022b) by interpolating the coordinates of x and y in a meshgrid format with a colour map that represented the neighbouring points. As depicted in Figure 1, it was observed that the clicks were predominantly concentrated in the heart and abdominal regions. The results of the best 5 emotional categories in the ranking were used to score the intensity of 33 emotions. That is, the first, second, third, fourth, and fifth categories were each scored as a 5, 4, 3, 2, and 1 point. The scores of each 33 emotional categories were then averaged for all participants (see Figure E in the Supplementary for all results).

We first performed the Shapiro–Wilk test for normality on the total number of clicks at cardiac and abdomen positions in each participant, the aesthetic scores of the multiple-choice categorical judgements, and the valence and arousal scores of the nine-point dimensional judgments. Depending on the result of the test for normality, either the parametric or non-parametric (Friedman) repeated-measure analyses of variance (ANOVA) were applied to compare the total number of clicks at cardiac and abdomen positions, and the scores of valences, arousals, and aesthetic appreciation, among different types of chord progressions. The dependent variable was the total number of clicks for each cardiac and abdomen area and the scores of valences, arousals, and aesthetic appreciation, and the within-subject factor was 8 types of chord progressions. Further, either the parametric or non-parametric (Spearman) correlation analysis was applied to understand how the total numbers of clicks at cardiac and abdomen positions were correlated with the scores of valences, arousals, and aesthetic appreciation. Statistical analyses were conducted using jamovi Version 1.2 (The jamovi project, 2021). We selected *p* < .05 as the threshold for statistical significance and used a false discovery rate method for multiple comparison testing.

Then, to visualize the difference in the emotional distributions between the different types of chord progressions and between groups who rated high and low valence, we compared the scores of the emotional distributions of 33 emotional category by Uniform Manifold Approximation and Projection (UMAP). For each of the 92 unique chord progressions encompassed within the eight types of chord progressions, the 33 emotional categories were averaged separately for groups with valence values above 6 and those below 4. Subsequently, using the averaged data from these 33 emotional categories, we employed UMAP in Python to craft a two-dimensional representation. Points in the plot were colour-coded based on the eight types of chord progressions. Furthermore, high-valence groups were represented by triangles, while low-valence groups were denoted with stars (⍰). Further, to visualize the difference in the emotional distributions between the different types of chord progressions and between groups who showed strong and weak sensations of each cardiac and abdomen area, we compared the scores of the emotional distributions of 33 emotional categories by UMAP. For each of the 92 unique chord progressions encompassed within the eight types of chord progressions, the 33 emotional categories were averaged separately for groups above 1 (i.e., felt bodily sensation. at cardiac or abdomen areas) and those 0 (i.e., non-bodily sensation. at cardiac or abdomen areas). Subsequently, using the averaged data from these 33 emotional categories, we employed UMAP in Python to craft a two-dimensional representation. Points in the plot were colour-coded based on the eight types of chord progressions. Furthermore, high-valence groups were represented by the solid circle, while low-valence groups were denoted with a cross mark (⍰). In all the UMAP analyses, the number of components was 2, the random state value was 42, the minimum distance value was 0.001, and the spread value was set to 10.

## Acknowledgements

This research was supported by the Japan Science and Technology Agency (JST) Moonshot Goal 9 (JPMJMS2296), Japan. The funding sources had no role in the decision to publish or prepare the manuscript.

